# *Yersinia pestis* genomes reveal plague in Britain 4,000 years ago

**DOI:** 10.1101/2022.01.26.477195

**Authors:** Pooja Swali, Rick Schulting, Alexandre Gilardet, Monica Kelly, Kyriaki Anastasiadou, Isabelle Glocke, Tony Audsley, Louise Loe, Teresa Fernández-Crespo, Javier Ordoño, David Walker, Tom Davy, Marina Silva, Mateja Hajdinjak, Anders Bergström, Thomas Booth, Pontus Skoglund

## Abstract

Extinct lineages of *Yersinia pestis*, the causative agent of the plague, have been identified in several individuals from Central Europe and Asia between 5,000 and 3,500 years before present (BP). One of these, the ‘LNBA lineage’ (Late Neolithic and Bronze Age), has been suggested to have spread into Central Europe with human groups expanding from the Eurasian steppes. Here, we show that LNBA plague was spread to Europe’s northwestern periphery by sequencing *Yersinia pestis* genomes from two individuals dating to ~4,000 cal BP from an unusual mass burial context in Somerset, England, UK. This represents the earliest evidence of plague in Britain documented to date. These British *Yersinia pestis* genomes belong to a sublineage previously observed in two Bronze Age individuals from Central Europe that had lost the putative virulence factor *yapC*. This sublineage is later found in Central Asia ~3,600 BP. While the severity of disease is currently unclear, the wide geographic distribution within a few centuries suggests substantial transmissibility.

## Introduction

*Yersinia pestis*, the causative agent of the plague, is a zoonotic bacteria which can be transmitted via the bite of an infected flea vector, causing either bubonic or septicemic plague, or via respiratory droplets through human-to-human contact causing pneumonic plague. Ancient DNA analysis has identified *Yersinia pestis* as the causative agent of not only the Late Neolithic and Bronze Age (LNBA) plague^1–5^, but also of historic pandemics such as the Justinianic plague^6–8^ and the Black Death^9,10^. The LNBA lineages lack the *ymt* virulence factor which enables survival of the bacteria in the midgut of fleas^1^, and are therefore likely to have been spread via the respiratory route. The first known evidence of the *ymt* gene dates to ~3,800 BP in Samara, Russia^4^. The LNBA lineage was likely brought to Europe in the Late Neolithic by human expansions originating in the Eurasian steppes^1,11,12^, and it has been hypothesised that this lineage contributed to the decline of certain Late Neolithic European societies^1,2^. However, it has been unclear how far the LNBA plague spread throughout Europe, with the most westward findings so far coming from present-day southern Germany ~3,500 BP^3^. The burials in Germany were associated with the Bell Beaker archaeological complex which linked continental Europe with Britain, opening the possibility that Bell Beaker groups from western Europe were also affected by outbreaks of plague, although this has yet to be observed directly. The extent to which plague spread across Bronze Age Europe is uncertain, due to the questions around virulence, but also heterogeneity in genetic ancestry of Bell Beaker burials across Europe^13^, suggesting variability in levels of human interaction and dynamics of any disease spread.

## Results

We identified *Yersinia pestis* in 2 out of 16 individuals which were screened for pathogen DNA from a mass burial assemblage of disarticulated human remains recovered from a natural shaft near Charterhouse Warren Farm in Somerset, England (**Figure 1A**). More than 40 men, women and children are represented at the site. Modifications found on a significant proportion of bones suggests that these individuals had been subject to fatal perimortem trauma and subsequent processing before disarticulated bones were deposited together in the shaft in what is likely to have been a single event, though archaeological work is still ongoing. A number of objects associated with the Beaker culture were also found^14^. The two sampled teeth derived from separate individuals, children aged 10 ± 3 yr and 12 ± 3 yr. The mandible associated with one of the teeth has been directly radiocarbon dated to 4,145–3,910 cal BP (95.4% confidence; OxA-37840: 3,685 ± 30 BP). This date is consistent with two previously published radiocarbon dates on human bones from this assemblage^15^.

**Figure 1.**
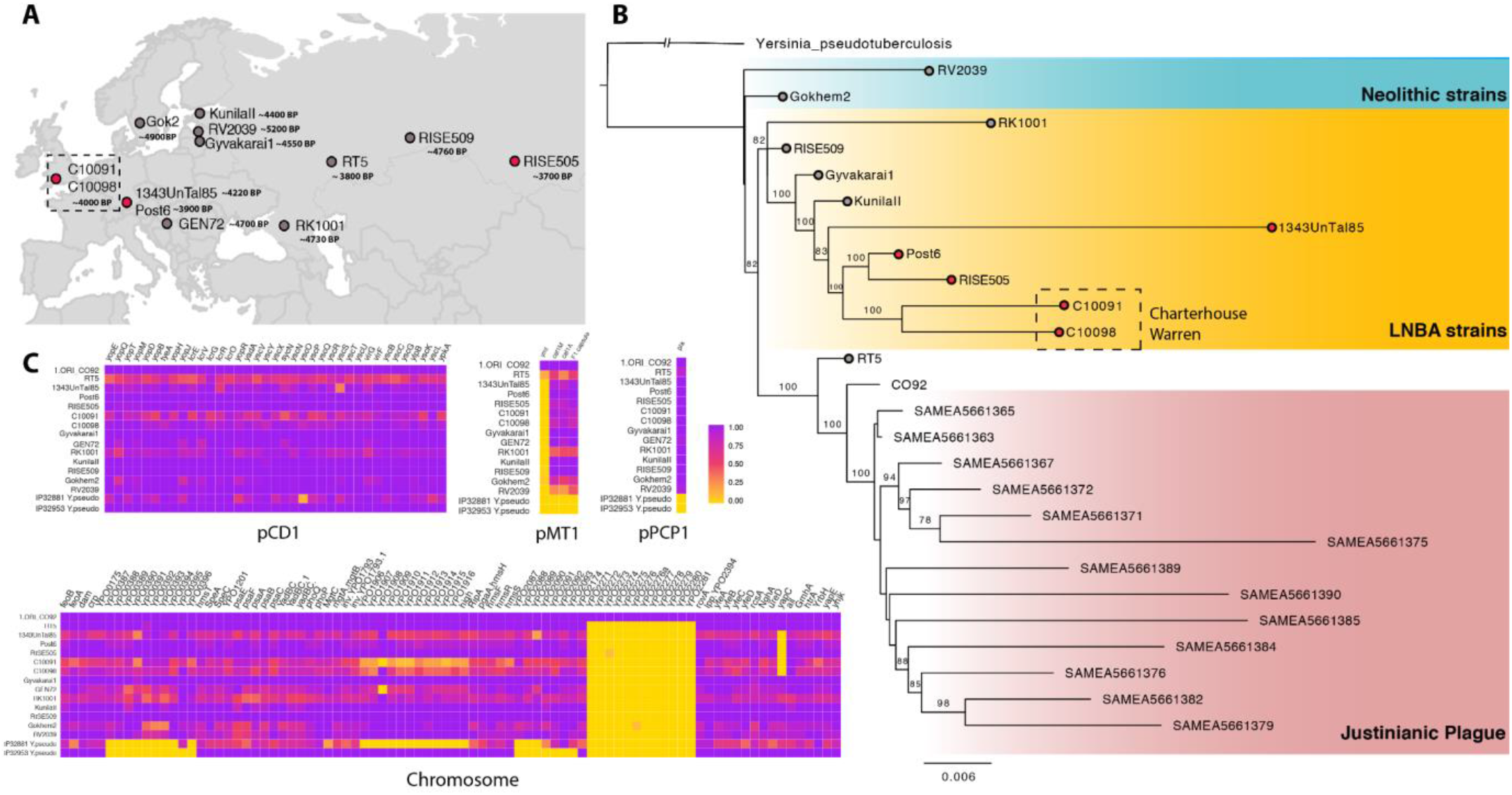
*Yersinia pestis* genomes from Bronze Age Britain. A) Map of prehistoric *Yersinia pestis* genomes. Genomes representing the monophyletic young LNBA lineages are highlighted in red. B) Phylogeny reconstructed with two Charterhouse Warren genomes and 23 previously published ancient *Yersinia pestis* genomes^1–8^ (branches with tip circles represented in Figure 1A, one modern-day genome (CO92) and two collapsed *Yersinia pseudotuberculosis* outgroups in RaXML. Neolithic lineages are highlighted in blue, LNBA lineages in yellow and Justinianic plague lineages in pink. Individuals from this study are outlined with a dashed box. Individuals with tip lables are also present in Figure 1A, with red tip lables representing the monopyhyletic young LNBA lineage. Node labels indicate bootstrap support values. C) Heatmap of inferred presence/absence of virulence assessed by depth-of-coverage across *Yersinia pestis* Chromosome and plasmids pCD1, pMT1 and pPCP1 (**Methods**).

### Ancient DNA retrieval

All samples were processed in a dedicated ancient DNA facility at the Francis Crick Institute. Dentine samples were taken from the cementum enamel junction and single-stranded DNA libraries were prepared from extracts made from the resulting powder^16^ (**Methods**). Among the individuals screened, we identified two samples (C10091 and C10098) that had an excess number of observed sequence matches to *Yersinia pestis* compared to *Yersinia pseudotuberculosis*. In addition, 764 million reads were obtained for individual C10098 with direct shotgun sequencing, after size selection to enrich for molecules at least 35 bp in length^16^ (**Methods**). Additionally, libraries underwent one round of hybridization with an in-solution target enrichment approach using *Yersinia pestis* RNA baits, resulting in an average 2-fold and 1.5-fold coverage of the *Yersinia pestis* genome for C10091 and C10098, respectively (**Table 1**). Both genomes showed evidence of authenticity by three central criteria: postmortem damage; the distribution of edit distances among sequences; and breadth of coverage across the genome (Supplementary **Figure 1**).

**Table 1.**
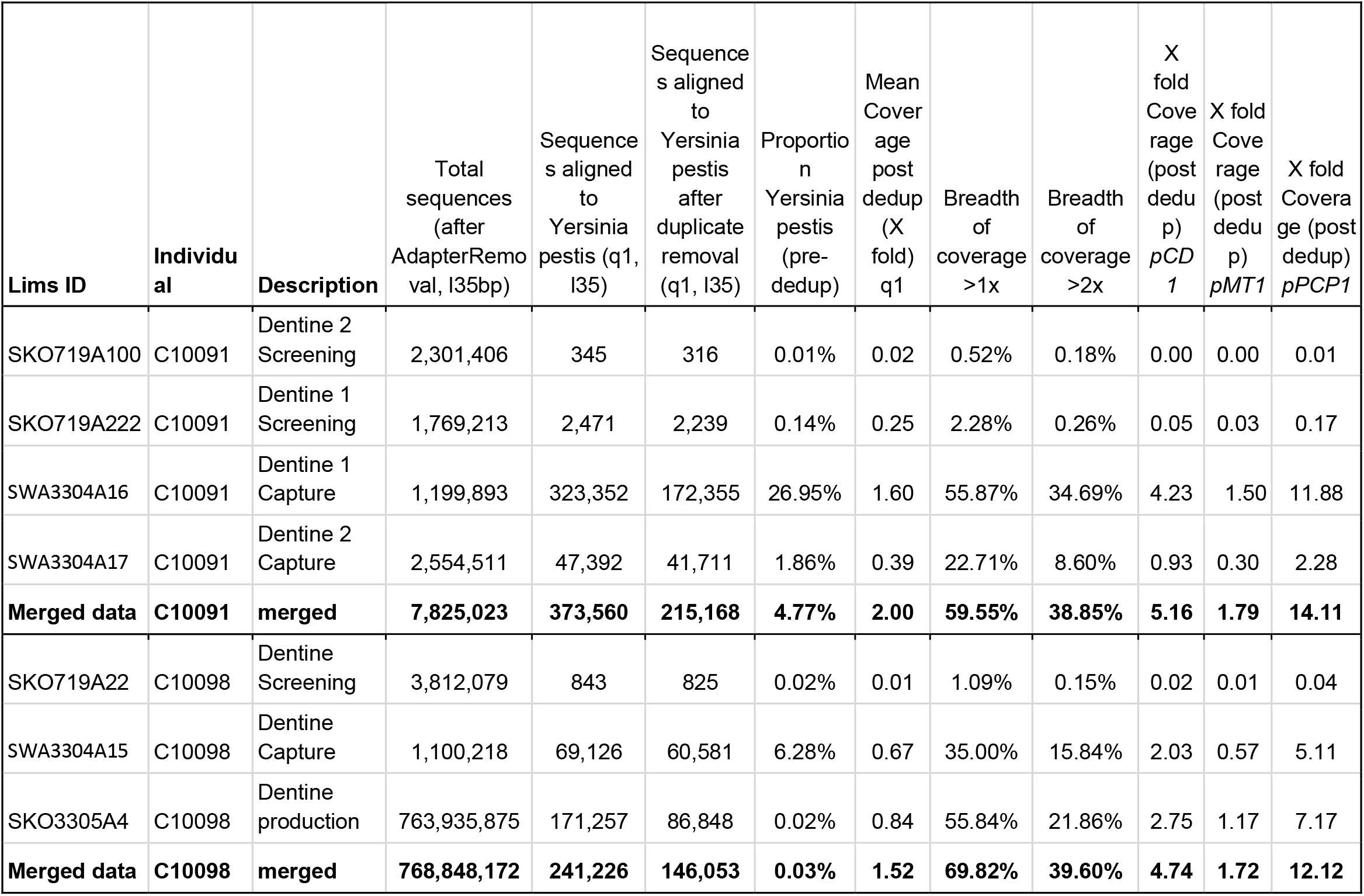
Sequencing statistics for different libraries and experiments

### Phylogenetic and functional analysis

In a phylogeny with 25 ancient and modern *Yersinia pestis* strains and *Yersinia pseudotuberculosis* outgroups, the Charterhouse Warren *Yersinia pestis* genomes form a clade with previously identified Bronze Age strains from Augsburg, Germany^3^ **(Figure 1B)**. A 34.2kb region found to be deleted in the youngest representatives of the LNBA lineages was also absent in the Charterhouse Warren genomes. The *yapC* virulence factor, which has been shown *in vitro* to mediate adhesion in cultured cells and could thus have enhanced colonisation of *Yersinia pestis* during infection^17^, and which was absent in the youngest LNBA genomes, was also absent in the Charterhouse Warren genomes (**Figure 1C**). A separate 36.4kb region that is absent in RISE505 (Central Russia), but present in Post6 (Germany), was also present in both C10091 and C10098 individuals^3^. As with all previous LNBA genomes, all virulence factors on the plasmids were present, with the exception of the *ymt* gene. *UreD*, associated with increased flea toxicity and *pde-2* which is involved in the downregulating mechanism for biofilm formation and *flhD*, the inactivation of which is associated with immune invasion, are all functional in the Charterhouse Warren genomes, which is congruent with previous LNBA genomes^4^ suggesting flea-mediated transmission was less likely. Additionally, we compared the two Charterhouse Warren *Yersinia pestis* genomes and identified three transversions SNPs between them with a minimum coverage of three; a synonymous mutation in the *tufA* gene, a non-synonymous mutation in the *pgm* gene and a mutation in the *rnpB gene* (**Methods**).

## Discussion

Previous individuals from Bronze Age and Neolithic Europe showing *Yersinia pestis* infections of the LNBA lineage all came from single or multiple rather than mass burials^1,11,12^. The LNBA *Yersinia pestis* genomes we present from Britain are the first to come from a mass burial context. Moreover, mass burials of disarticulated human remains in caves are rare in Bronze Age Britain and could represent a non-normative form of treatment. However, the striking evidence for fatal trauma amongst the Charterhouse Warren assemblage^14^ makes it unlikely that this mass burial was due to a deadly outbreak of plague.

Considered alongside other available genomes, our results show that in the period 4,100-3,800 BP *Yersinia pestis* was widespread from Britain in the West, to Central Asia in the East. The earliest known *Yersinia pestis* strain carrying the *ymt* gene has been found in Central Asia and dates to 3800 BP^4^. This is close in time to our two *Yersinia* genomes from Britain, providing further evidence for differential contemporary frequencies of the *ymt* gene in Europe and Central Asia. *Yersinia pestis* lineages in Britain had close affinities with Bronze Age genomes in present-day Germany, and a later lineage in Asia (RISE505) which may have arrived from Europe with Srubnaya-associated expansions^3^. Our results reveal that LNBA *Yersinia pestis* lineages were not confined to Bronze Age Central Europe, but were spread westward to Britain, and were thus present in the western Bell Beaker archaeological horizon.

## Methods

### Archaeological context

Charterhouse Warren is a natural shaft in the limestone of the Mendip Hills, Somerset. Initial excavations in the 1970s were instigated in order to determine whether the shaft led to a cave system, as no archaeological remains from the site were known at that time^14^. The assemblage of human remains sampled for aDNA was encountered in a distinct layer at ca. 15m depth, together with faunal remains and a small number of artefacts, including sherds of a Beaker vessel. At least 40 men, women and children are represented by fragmentary and disarticulated remains. Culturally and chronologically, the assemblage dates to the late Beaker period/Early Bronze Age, ca. 2200-2000 cal BC (ca. 4150-3950 cal BP). This is a non-normative burial context for this period, which is dominated by articulated burials under round barrows. While disarticulated remains are sometimes encountered, the scale of deposition at Charterhouse Warren is unique in a British context, suggesting that a very unusual event is represented. Modelling of the radiocarbon dates is ongoing, but it is possible that the remains were deposited as a single event. The evidence for blunt force perimortem trauma to a number of crania together with evidence for dismemberment suggests that this may relate to an episode of extreme violence.

### Sampling, DNA extraction and library preparation

Samples were processed at the Francis Crick Institute in a dedicated cleanroom facility. We used a EV410-230 EMAX Evolution Dentistry drill to clean the surface of the teeth and sampled both the cementum and multiple fractions of the dentine, resulting in 7-25mg of powder from the dentine. Dentine powders were then lysed with 300ul (<10mg of powder) or 1000ul (>10mg of powder) of lysis buffer (0.5 EDTA pH 8.0, 0.05% Tween-20, 0.25mg/ml Proteinase K^18^) and incubated overnight at 37°C. Lysates were centrifuged at 2 min at maximum speed (13,200 rpm) in a table centrifuge and 140ul of the lysate was transferred into FluidX tubes for automated extraction on an Agilent Bravo Workstation^19^. Extracts were turned into single-stranded DNA libraries^16^ (with no treatment to remove uracils), and were then double-indexed^20^ and underwent paired-end sequencing on the HiSeq4000 to 5 million reads per sample for initial screening. All samples were processed alongside negative lysate and extraction controls as well as positive and negative library controls.

### Target enrichment

Following initial pathogen detection, libraries were taken forward for target enrichment using *Yersinia pestis* baits designed by myBaits Arbor Biosciences, following myBaits custom RNA seq v5.1 (March 2021) High Sensitivity protocol^21^. We used 7ul of the initial library for one round of hybridization at 55°C with 23-hour overnight incubation and 20 PCR cycles, followed by a heteroduplex removal using a one cycle PCR and MinElute Purification (Qiagen) clean up. These enriched libraries were sequenced to ~3 million reads per library on the Illumina MiSeq.

### Size selection for shotgun sequencing

Before production sequencing, fragments smaller than 35bp and larger than 150bp were removed from the library, as in Gansauge et al. 2020^16^. Specifically, 100ng of the initial library were biotinylated and streptavidin beads were used to isolate the non-biotinylated strand and obtain a single-stranded library. This sample (pooled with 3 others) was then loaded on a denaturing polyacrylamide gel along with 35bp and 150bp insert markers and fragments within the desired sequence length were physically excised and eluted from the gel, after an overnight incubation. The resulting size-selected library was further amplified and 764 million reads were sequenced on the Illumina NovaSeq.

### Bioinformatic processing

Samples were processed via the nf-core/eager v2 pipeline^22^. First, adapters were removed, paired-end reads were merged and bases with a quality below 20 were trimmed using AdapterRemoval v2^23^ with –trimns –trimqualities –collapse –minadapteroverlap 1 and –preserve5p. Merged reads with a minimum length of 35bp were mapped to the hs37d5 human reference genome with Burrows-Wheeler Aligner (BWA-0.7.17 aln)^24^ using the following parameters “-l 16500 -n 0.01 -o 2 -t 1” ^25,26^, and unmapped reads were then processed in Kraken2^27^ to detect k-mers matching *Yersinia pestis*.

Sequence data from capture and shotgun libraries were merged and aligned to the CO92 *Yersinia pestis* reference genome (NC_003143.1) and the CO92 plasmids, pMT1 (NC_003134.1), pCD1 (NC_003131.1) and pPCP1 (NC_003132.1) with BWA aln^24^ with parameters as above. Duplicates were removed by keeping only the first sequence out of any set of sequences with the same start position and length (https://github.com/pontussk/samremovedup). Aligned BAM files were authenticated using the following three criteria^28^: the observation of postmortem damage via *PMDtools*^29^, a majority of reads showing an edit distance of 1 or less from the reference genome, and an even breadth of coverage across the reference genome (**Figure S1**).

### Analysis of functional elements and virulence factors

We used previously identified virulence genes on the *Yersinia pestis* chromosome, pCD1, pMT1 and pPCP1 plasmids, and analysed the coverage of these genes with BEDTools v2.29.2 (Quinlan & Hall 2010) (**Figure 1C**). The ggplot2^30^ package in R was then used to make a heatmap of the percentage of positions in each virulence gene covered by at least one sequence. Additionally, we inspected previously identified indels to check whether they were present or absent in C10091 and C10098. Comparison of both Charterhouse strains was done using HTSbox (http://bit.ly/HTSBox) pileup and filtering for minimum coverage of 3 sequences per site on both strains, excluding all transitions.

### Phylogenetic reconstruction

External data from Bronze age and Neolithic plague genomes were downloaded from the European Nucleotide Archive (ENA) and processed using the same parameters as data generated in this study, described above (**Bioinformatic processing**). Additionally, two strains of *Yersinia pseudotuberculosis* (IP32881 and IP32953) were aligned to the CO92 reference genome. For IP32953 which was in a fasta format, MiniMap2-2.17^31^ was used.

After duplicate read removal, BAMs were filtered and converted to majority-call (i.e. the base that is observed in the majority of overlapping reads is taken as the consensus) fasta format via HTSbox (https://github.com/lh3/htsbox) pileup using the following parameters: -M -q 30 -Q 30 -l 35, with a minimum majority base count of -s 2 for ancient genomes and -s 1 for modern-day genomes. We then filtered these fasta files to keep only polymorphic sites with transversions only, exclude individuals with more than 70% missingness and removing sites that were missing in more than 70% of the remaining individuals, resulting in 13,474 sites that were taken forward to RAxML^32^. We used a GTRGAMMA substitution model with rapid bootstrapping parameters, with 100 bootstraps; The maximum likelihood phylogeny was rooted in FigTree^33^, and the two *Yersinia pseudotuberculosis* outgroups (IP32881 and IP32953) were collapsed for the final display in **Figure 1B**.

## Acknowledgements

We thank the Advanced Sequencing Facility at the Francis Crick Institute for technical support. This work was supported by the Vallee Foundation, the European Research Council (grant no. 852558), the European Molecular Biology Organisation, the Wellcome Trust (217223/Z/19/Z), and Francis Crick Institute core funding (FC001595) from Cancer Research UK, the UK Medical Research Council, and the Wellcome Trust. M.H. was supported by Marie Skłodowska Curie Actions (grant no. 844014). Osteological analyses of Charterhouse Warren human skeletal assemblage were supported by the British Academy (SG163375), with funding for radiocarbon dating supplied through NERC’s NEIF programme (NF/2018/1/3). This research was funded in whole, or in part, by the Wellcome Trust (FC001595 and 217223/Z/19/Z). For the purpose of Open Access, the author has applied a CC BY public copyright licence to any Author Accepted Manuscript version arising from this submission.

## Author contributions

Conceptualization: P. Sw., P. Sk

Data curation: P. Sw., A. G.

Formal Analysis: P. Sw.

Funding acquisition: R. S., P. Sk.

Investigation: P. Sw., R. S., A. G., M.K, K.A., I.G., T.D., M.S., M.H., A.B., T. B., P.Sk.

Project administration: P.Sw., R.S., T.B., P.Sk.

Resources: R. S., T.A., L.L., L.L., TFC, J.O.

Supervision: P. Sk.

Visualization: P. Sw., A.G., K.A.

Writing – original draft: P. Sw., P. Sk.

Writing – review & editing: P. Sw., R. S., M.K., K. A., M.H., A.B., T.B., P. Sk.

## Data availability

All sequence data will be available in the European Nucleotide Archive upon publication.

## References

1. Rasmussen, S. et al. Early divergent strains of Yersinia pestis in Eurasia 5,000 years ago. Cell 163, 571–582 (2015).

2. Rascovan, N. et al. Emergence and Spread of Basal Lineages of Yersinia pestis during the Neolithic Decline. Cell 176, 295–305.e10 (2019).

3. Andrades Valtueña, A. et al. The Stone Age Plague and Its Persistence in Eurasia. Curr. Biol. 27, 3683–3691.e8 (2017).

4. Spyrou, M. A. et al. Analysis of 3800-year-old Yersinia pestis genomes suggests Bronze Age origin for bubonic plague. Nat. Commun. 9, 2234 (2018).

5. Susat, J. et al. A 5,000-year-old hunter-gatherer already plagued by Yersinia pestis. Cell Rep. 35, 109278 (2021).

6. Harbeck, M. et al. Yersinia pestis DNA from skeletal remains from the 6(th) century AD reveals insights into Justinianic Plague. PLoS Pathog. 9, e1003349 (2013).

7. Keller, M. et al. Ancient Yersinia pestis genomes from across Western Europe reveal early diversification during the First Pandemic (541-750). Proc. Natl. Acad. Sci. U. S. A. 116, 12363–12372 (2019).

8. Feldman, M. et al. A High-Coverage Yersinia pestis Genome from a Sixth-Century Justinianic Plague Victim. Mol. Biol. Evol. 33, 2911–2923 (2016).

9. Bos, K. I. et al. A draft genome of Yersinia pestis from victims of the Black Death. Nature 478, 506 (2011).

10. Spyrou, M. A. et al. Historical Y. pestis Genomes Reveal the European Black Death as the Source of Ancient and Modern Plague Pandemics. Cell Host Microbe 19, 874–881 (2016).

11. Haak, W. et al. Massive migration from the steppe was a source for Indo-European languages in Europe. Nature 522, 207 (2015).

12. Allentoft, M. E. et al. Population genomics of Bronze Age Eurasia. Nature 522, 167–172 (2015).

13. Olalde, I. et al. The Beaker phenomenon and the genomic transformation of northwest Europe. Nature 555, 190–196 (2018).

14. Levitan, B.M., Audsley, A., Hawkes, C.J.L., B.M., Moody, A.A.D., Moody, P.D., Smart, P.L. and Thomas, J.S. Charterhouse Warren Farm Swallet, Mendip, Somerset: Exploration, Geomorphology, Taphonomy and Archaeology: Proceedings University of Bristol Spelaeological Society 18(2) 1988 P171-239. Proceedings of the University of Bristol Spelaeological Society 171–239.

15. Levitan, B.M and Smart, P.L. Charterhouse Warren Farm Swallet, Mendip, radiocarbon dating evidence. Proceedings of the University of Bristol Spelaeological Society 18, 390–394 (1989).

16. Gansauge, M.-T., Aximu-Petri, A., Nagel, S. & Meyer, M. Manual and automated preparation of single-stranded DNA libraries for the sequencing of DNA from ancientbiological remains and other sources of highly degraded DNA. Nat. Protoc. 15, 2279–2300 (2020).

17. Felek, S., Lawrenz, M. B. & Krukonis, E. S. The Yersinia pestis autotransporter YapC mediates host cell binding, autoaggregation and biofilm formation. Microbiology 154, 1802–1812 (2008).

18. Dabney, J. et al. Complete mitochondrial genome sequence of a Middle Pleistocene cave bear reconstructed from ultrashort DNA fragments. Proc. Natl. Acad. Sci. U. S. A. 110, 15758–15763 (2013).

19. Rohland, N., Glocke, I., Aximu-Petri, A. & Meyer, M. Extraction of highly degraded DNA from ancient bones, teeth and sediments for high-throughput sequencing. Nat. Protoc. 13, 2447–2461 (2018).

20. Kircher, M., Sawyer, S. & Meyer, M. Double indexing overcomes inaccuracies in multiplex sequencing on the Illumina platform. Nucleic Acids Res. gkr771 (2011).

21. Furtwángler, A. et al. Comparison of target enrichment strategies for ancient pathogen DNA. Biotechniques 69, 455–459 (2020).

22. Yates, J. A. F. et al. Reproducible, portable, and efficient ancient genome reconstruction with nf-core/eager. doi:10.1101/2020.06.11.145615.

23. Schubert, M., Lindgreen, S. & Orlando, L. AdapterRemoval v2: rapid adapter trimming, identification, and read merging. BMC Res. Notes 9, 88 (2016).

24. Li, H. & Durbin, R. Fast and accurate short read alignment with Burrows–Wheeler transform. Bioinformatics 25, 1754–1760 (2009).

25. Meyer, M. et al. A High-Coverage Genome Sequence from an Archaic Denisovan Individual. Science 338, 222–226 (2012).

26. Poullet, M. & Orlando, L. Assessing DNA Sequence Alignment Methods for Characterizing Ancient Genomes and Methylomes. Frontiers in Ecology and Evolution 8, 105 (2020).

27. Wood, D. E., Lu, J. & Langmead, B. Improved metagenomic analysis with Kraken 2. Genome Biol. 20, 257 (2019).

28. Key, F. M., Posth, C., Krause, J., Herbig, A. & Bos, K. I. Mining Metagenomic Data Sets for Ancient DNA: Recommended Protocols for Authentication. Trends Genet. 33, 508–520 (2017).

29. Skoglund, P. et al. Separating endogenous ancient DNA from modern day contamination in a Siberian Neandertal. Proceedings of the National Academy of Sciences (2014) doi:10.1073/pnas.1318934111.

30. Wickham, H. ggplot2: Elegant Graphics for Data Analysis. (Springer, New York, NY, 2009).

31. Li, H. New strategies to improve minimap2 alignment accuracy. Bioinformatics (2021) doi:10.1093/bioinformatics/btab705.

32. Stamatakis, A. RAxML version 8: a tool for phylogenetic analysis and post-analysis of large phylogenies. Bioinformatics 30, 1312–1313 (2014).

33. Rambaut, A. FigTree v1.3.1: Tree Figure Drawing Tool (Available from http://tree.bio.ed.ac.uk/software/figtree/) Accessed November 2, 2011. (2009).

