## Supplementary Figure 1 for "*Yersinia pestis* genomes reveal plague in Britain 4,000 years ago"

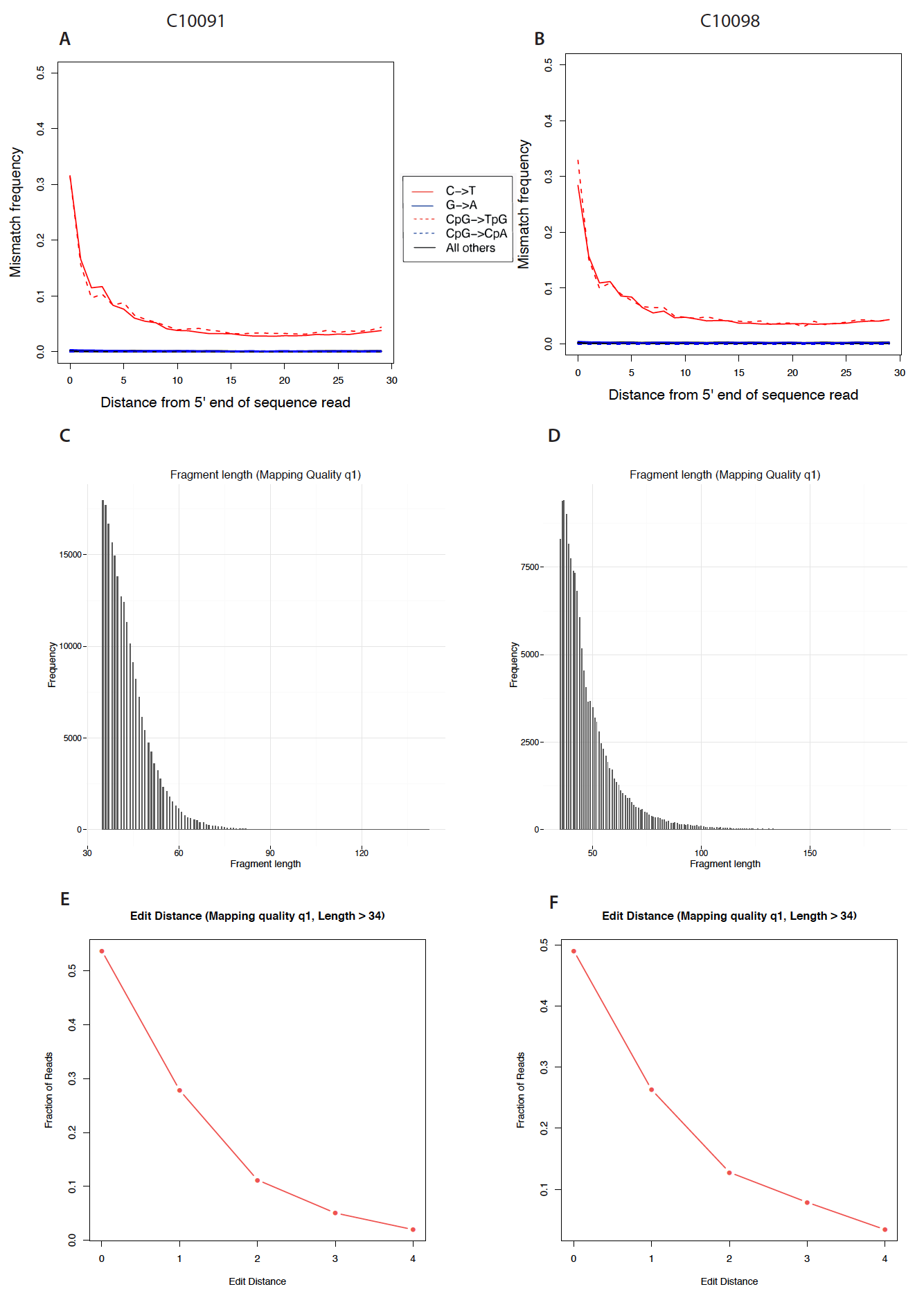

**Supplementary Figure 1**. **Authentication metrics for *Yersinia pestis* genomes from Charterhouse Warren.**

A-B) Nucleotide misincorporations resulting from cytosine deamination. C-D) Fragment length distribution.E-F) Number of sequences with edit distance of 1 to 4 from the *Yersinia pestis* reference.
